# Competition and interdependence define multifaceted interactions of symbiotic *Nostoc* sp. and *Agrobacterium* sp. under inorganic carbon limitation

**DOI:** 10.1101/2024.07.16.603663

**Authors:** Jonna E. Teikari, David A. Russo, Markus Heuser, Otto Baumann, Julie A. Z. Zedler, Anton Liaimer, Elke Dittmann

## Abstract

Cyanobacteria of the *Nostoc* genus are capable of forming symbiotic relationships with plants, thus transitioning to a heterotrophic lifestyle in return for providing bioavailable nitrogen to the host. The diazotrophic photoautotrophs also serve as a hub for specialized heterotrophic bacterial communities whose physiological contributions are poorly understood. By comparing the axenic strain *Nostoc punctiforme* PCC 73102 and the related strains *Nostoc* sp. KVJ2 and KVJ3, which still maintain their heterotrophic microbiome, we were able to demonstrate an almost obligate dependence of the cyanobacteria on the heterotrophic partners under carbon-limiting conditions. Detailed analysis of the intimate bilateral relationship between *N. punctiforme* and the isolate *Agrobacterium tumefaciens* Het4 using shotgun proteomics and microscopy uncovered a complex partnership characterized, among other traits, by competition for iron and facilitation for carbon. Although competitive interactions with *A. tumefaciens* Het4 compromise nitrogen fixation and stimulate the degradation of cyanophycin, mutualistic dependency prevails under inorganic carbon limitation. Both the absence of the high affinity bicarbonate uptake transporter SbtA and the prevalent extracarboxysomal localization of the carbon-fixing enzyme RubisCO, as detected by immunofluorescence microscopy, suggest that a weak carbon concentrating mechanism in *N. punctiforme* enforces a dependence on heterotrophic bacteria. Further, immunofluorescence, electron microscopic and proteomic analyses reveal a pronounced extracellular recycling of proteins under N- and C-limiting conditions. Our study shows that the pivotal influence of heterotrophic bacteria on symbiotic *Nostoc* strains should be considered when analyzing these cyanobacteria, especially in the free-living state. This work also sheds new light on how *Nostoc* benefits from the organic carbon provided by plant hosts.

## Introduction

Cyanobacteria are capable of maintaining a wide range of symbiotic relationships with eukaryotic hosts such as plants, fungi and protists (Rai et al., 2000). Plant-associated *Nostoc* species play an essential role in terrestrial ecosystems, most importantly due to their ability to fix atmospheric nitrogen and further fuel the ecosystem by transferring newly assimilated nitrogen directly to their host plant (Rousk et al., 2013). In such epiphytic or endophytic symbioses, *Nostoc* releases most of the fixed nitrogen, mainly in the form of ammonia, to the host (Meeks, 1998; DeLuca et al., 2008; Larmola et al., 2014). In return, the plant provides the cyanobacteria shelter and, more importantly, organic carbon (Chapman and Margulis, 1998).

The ecology of the terrestrial and symbiotic *Nostoc* is generally well characterized. A complex lifestyle is maintained by large genomes that encode structures and physiological traits for specialized cells: nitrogen-fixing heterocysts, resting cells, akinetes and motile filaments, known as hormogonia, which have a great importance in the formation of the plant symbiosis (Meeks and Elhai, 2002; Risser, 2023). *Nostoc* are also a rich source of bioactive natural products that have versatile but partially unknown roles in inter- and intraspecific communication within their ecosystem (Liaimer, 2011; Liaimer et al., 2015; D’Agostino, 2023) .

Although cyanobacteria are photoautotrophic organisms, and thus generally considered independent of organic carbon sources, many of them are, in fact, facultative heterotrophs and dependent on the organic carbon supply from their ecosystem (Pelroy et al., 1972; Yu et al., 2009). In symbiosis with plants they take advantage of a heterotrophic lifestyle (Adams and Duggan, 2008) to alleviate more energy for nitrogen fixation. Organic carbon is typically transported in small molecules such as glucose, but recent findings have shown that symbiotic *Nostoc* sp. have acquired a set of beneficial genes to transport and degrade more complex carbon sources, such as cell wall components (Khamar et al., 2010; Warshan et al., 2017). The accumulation of beneficial genes over the course of evolution suggests plasticity in the carbon acquisition strategies, and the ability to utilize complex carbon sources may increase the fitness of cyanobacteria in plant symbiosis (Warshan et al., 2017; Warshan et al., 2018b).

In contrast, free-living *Nostoc* colonies, sometimes forming biofilms that may reach centimeters in diameter, rely on their own photosynthetic capacity and the availability of atmospheric CO_2_ which may become scarce in dense biofilms and large colonies. To cope better in low CO_2_ conditions, cyanobacteria have developed an effective carbon concentrating mechanism (CCM), that elevates CO_2_ near carboxysome-encapsulated RubisCO, the key enzyme involved in carbon fixation (Burnap et al., 2015). In marine and freshwater ecosystems, which are often C-limited *in situ* (Hein and SandJensen, 1997; Wilhelm et al., 2020), CO_2_ is also provided in exchange for dissolved organic carbon by the respiratory activities of heterotrophic bacteria tightly associated with cyanobacteria (Lee et al., 2017). *Nostoc*-associated microbiomes have so far been studied for *Nostoc flagelliforme* and *Nostoc commune,* which form gelatinous colonies in soil, and for spherical *Nostoc* colonies in freshwater ecosystems (Yue et al., 2016; Aguilar et al., 2019). While a pronounced habitat specificity was observed, the functional role of the bacteria in the *Nostoc* cyanosphere was not explicitly addressed. Therefore, the extent to which heterotrophic bacteria modulate the physiology of diazotrophic cyanobacteria remains largely unstudied. Notably, symbiotic *Nostoc* strains are difficult, sometimes impossible, to maintain as axenic isolates indicating intricate dependencies which are not yet understood (Heck et al., 2016; Warshan et al., 2018a).

Here we specifically compared the influence of the heterotrophic microbiome on the growth of *Nostoc* at different inorganic carbon concentrations. We selected three symbiotic *Nostoc* species: axenic *Nostoc punctiforme* PCC 73102, and non-axenic *Nostoc* sp. KVJ2 and *Nostoc* sp. KVJ3 (Liaimer et al., 2016). Despite having a similar genetic repository for carbon acquisition, axenic cyanobacteria were unable to proliferate under inorganic carbon limitation, while non-axenic cyanobacteria thrived.

Systematic investigation of the interaction between *N. punctiforme* PCC 73102 and the representative isolate *Agrobacterium tumefaciens* Het4 revealed a multifaceted relationship combining competition and facilitation for nutrients. The comparative analysis also provided insights into the weak carbon concentrating mechanism of *N. punctiforme* which could foster the dependency relationships between the partners. Our study sheds new light on the limited autonomy of *Nostoc* strains and highlights the importance of the heterotrophic bacterial community for maintaining a vital ecosystem.

## Results

### Heterotrophic microbiome drastically enhances the vitality of N. punctiforme PCC 73102

Three *Nostoc* strains PCC 73102, KVJ2 and KVJ3 were grown on plates of BG11_0_ medium without a nitrogen source containing low, medium, or high carbonate concentrations. In case of low carbon conditions, Na_2_CO_3_ was omitted while medium concentrations correspond to the standard amount of Na_2_CO_3_ found in BG11_0_. For high carbonate conditions, Na_2_CO_3_ was added tenfold the amount of standard BG11_0_ medium. The growth was followed for eight days on three independent replicate plates. Low and medium carbonate concentrations did not promote the growth of PCC 73102, whereas the highest availability of carbonate enabled fast growth of the strain (Figure 1A). In contrast to axenic PCC 73102, strains KVJ2 and KVJ3, both carrying the associated microbiomes, grew surprisingly well even at low carbonate levels, and showed virtually no difference in growth between limiting and high carbonate concentrations. The dependence of PCC 73102 on very high carbonate concentrations indicated a weak CCM of the strain (Figure. 1A).

**Figure 1.**
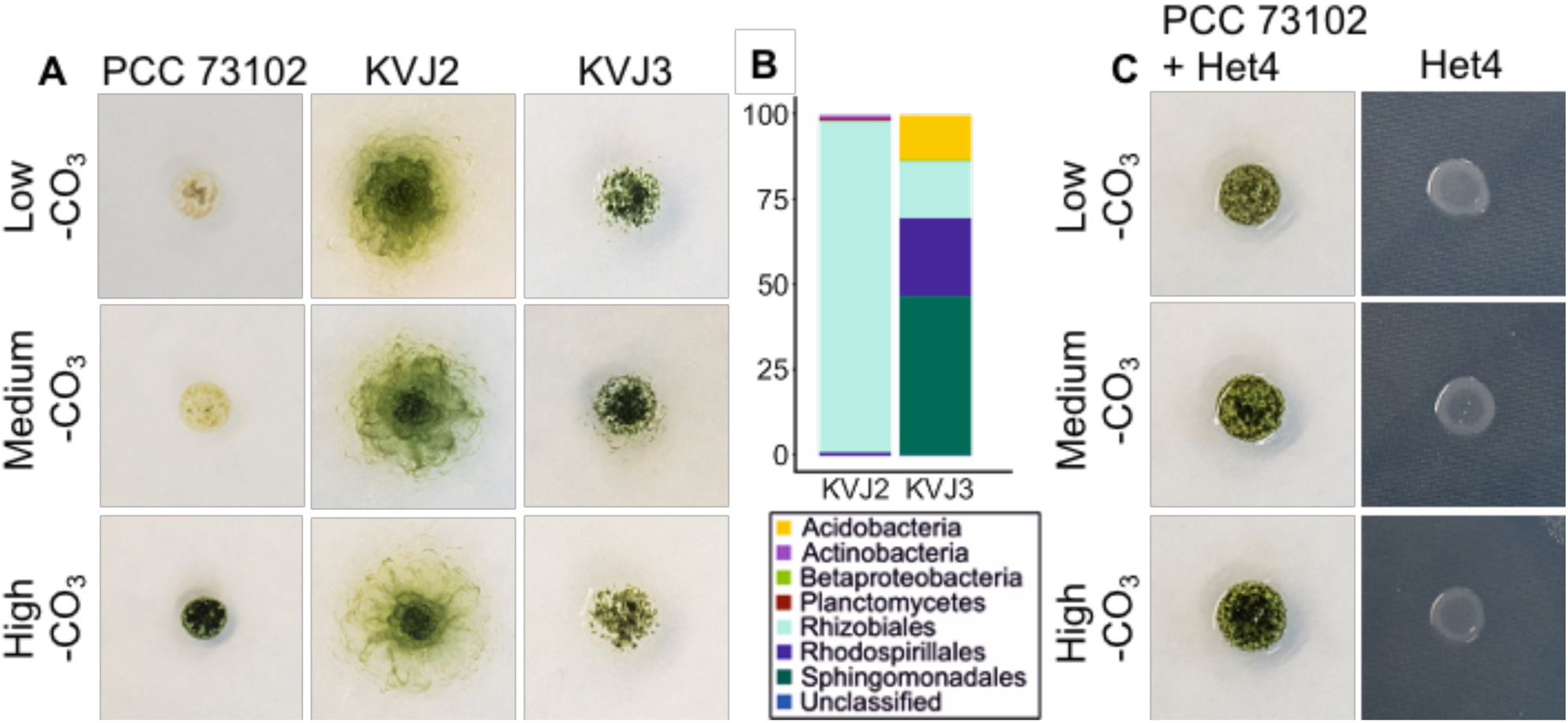
Growth of *Nostoc* sp. cyanobacteria is promoted by associated heterotrophic bacteria, particularly under inorganic carbon-limited conditions. (A) Axenic *N. punctiforme* PCC 73102 was unable to grow under low carbonate concentrations, whereas *Nostoc* sp. KVJ3 and KVJ2 thrived. (B) Microbiome analysis using 16S rRNA amplicon sequencing revealed a diversity of heterotrophic bacteria in *Nostoc* sp. KVJ2 and KVJ3 which were dominated by Alphaproteobacteria. (C) The isolate *A.tumefaciens* Het4 promotes growth of *N. punctiforme* PCC 73102 under carbon-limited conditions. See figure S3 for growth curves and digital image analysis.

To evaluate the adaptation strategies of the three studied *Nostoc* strains to niches with different inorganic carbon concentrations, the genetic repertoire of CCM-related genes was examined (Shibata et al., 2001; So and Espie, 2005; Sandrini et al., 2014). While the genome sequences of strain PCC 73102 and KVJ2 were publicly available in databases (GCA_000020025.1 and SAMN07173937), the sequence of strain KVJ3 was determined in the course of this study (PRJNA599284). All strains encoded the complete ATP-dependent bicarbonate uptake system BCT1 and BicA, along with NAD(P)H dehydrogenase complexes (Table S1), which are important for the proper functioning of the bicarbonate uptake system (Shibata et al., 2001). However, the high-affinity bicarbonate uptake transporter SbtA and the associated SbtB protein were absent from the genomes of all studied strains, suggesting a poor capability to cope with low levels of inorganic carbon, similar to the bloom-forming cyanobacterium *Microcystis* (Sandrini et al., 2014). Since all strains harbored the same set of genes encoding high- and low-affinity bicarbonate transporters, the vitality of the studied *Nostoc* cyanobacteria under inorganic carbon deficiency and their readiness to initiate plant symbiosis cannot be explained by a different genetic repertoire of CCM-related genes.

The genetic similarity in carbon acquisition and assimilation led us to anticipate that the heterotrophic bacterial communities play an important role in the vitality of free-living cyanobacteria under carbon limitation by providing CO_2_ and potentially other metabolites to the associated cyanobacteria. Thus, we studied the microbiomes of non-axenic strains KVJ2 and KVJ3 in more detail. Qualitative sequencing of 16S rRNA gene amplicons spanning the V3-V4 regions showed that both strains were heavily colonized by Proteobacteria (Figure 1B, Figure S1). The microbiome of KVJ2 was more consistent, mainly comprising Rhizobiales, while the microbiome of KVJ3 had greater diversity, with additional presence of Sphingomonadales, Rhodospirillales and Acidobacteria.

Four different colony types of heterotrophic bacteria monocultures were isolated from KVJ3 (Het1-4) and three from KVJ2 (Het5-7) using R2A agar plates. Phylogenetic analysis of 16S rRNA gene sequences revealed that four isolates (Het1, 2, 4, 5 and 6) belonged to the *Rhizobium*/*Agrobacterium* group (Alphaproteobacteria), with Het1 and Het2 representing the same species (Figure S2). Thus, Het2 was omitted from further analysis. Het3 represented *Achromobacter* (Betaproteobacteria) genus whereas *Paenibacillus* sp. Het7, belonging to Firmicutes, was the only isolate beyond Proteobacteria.

The isolation method used here showed clear selectivity towards *Rhizobium/Agrobacterium* and omitted Sphingomonadales and Acidobacteria, even though their abundance was evident based on the 16S rRNA amplicon metagenomic data. Notably, closely related strains of the *Rhizobium*/*Agrobacterium* group were also isolated after incidental laboratory contamination of the axenic strain PCC 73102 and isolated strains were included in the phylogenetic analysis as strains Het8 and Het9 (Figure S1).

Next, we tested the growth-promoting effects of the isolated heterotrophic bacteria on PCC 73102 by mixing axenic PCC 73102 with each of the isolates under BG11_0_ conditions. The growth was followed for eight days at medium carbonate concentration. Isolates Het1, 3, 4 and 8 strongly supported the growth of the cyanobacterium, and Het7 supported growth to some extent (Figure S2A). In co-cultures with Het5 and Het6, axenic cyanobacteria did not show signs of elevated growth, which points towards a great functional diversity of associated heterotrophic bacteria of the *Rhizobium*/*Agrobacterium* group.

The bacterial isolate *A. tumefaciens* Het4 (hereafter Het4) was selected to further understand the benefits of the microbial community for the cyanobacterium. This choice was based on the ability of Het4 to maintain physical interaction with PCC 73102 (Figure S2B) and to strongly enhance the growth of the cyanobacterium. The impact of the Het4 co-cultivation on growth of PCC 73102 was analyzed in replicate experiments for low, medium and high carbonate concentration and quantified by digital image analysis. A pronounced growth promotion effect was particularly evident under medium carbonate conditions in BG11_0_ (Figure 1C and S3). In contrast, the opposite effect was found at high carbonate concentrations. While PCC 73102 grew faster under these conditions, the Het4 interaction had a negative effect on growth. These reciprocal growth effects indicate that Het4 can counteract C-limitation in particular, but that the interaction is not solely mutualistic (Figure S3). Next, we sequenced the genome of isolate Het4. Analysis of amino acid identity (AAI) showed closest relatedness of the newly isolated strain to the *Agrobacterium* complex, which includes the well-known plant pathogen *Agrobacterium tumefaciens* str. C58 (Figure S4). Similar to strain C58 (Wood et al., 2001), the recently sequenced Het4 consisted of a circular chromosome (2.9 Mbp), a linear chromosome (2.1 Mbp), a cryptic plasmid (0.4 Mbp) and a tumor-inducing plasmid (0.37 Mbp). Het4 showed a very broad potential for the utilization of sugars, sugar alcohols and aromatic compounds as described for the reference strain C58. Further, Het4 carried the *nos* operon, which is required for the final step of the full denitrification pathway, and C58 is capable to directly ammonify nitrate and nitrite. Neither Het4 nor C58 had known pathways for nitrogen or carbon fixation.

### Heterotrophic bacteria modulate the physiology of N. punctiforme PCC 73102

To gain a better understanding of the interaction between Het4 and the cyanobacterium, the PCC 73102 proteome was compared in the presence and absence of Het4 using a shotgun proteomic approach. To this end, samples for endoproteome analysis were collected after eight days of cultivation on BG11_0_ agar plates with and without Het4. In total, 4078 proteins were identified in the PCC 73102 monoculture and 4062 were identified in co-culture with Het4. 4023 proteins were identified in both conditions and, in the presence of Het4, 123 proteins were significantly upregulated and 117 proteins were downregulated (FC ≤ -2 & ≥ 2 and adjusted p-value < 0.05) (Figure S5 and Dataset S1). Remarkably, amongst the set of upregulated proteins, 17 phycobilisome subunits were identified (Figure S5 and Dataset S1). This supports the morphological differences observed in mono- and co-culture where PCC 73102 exhibits strong bleaching when grown in low/medium inorganic carbon concentrations in the absence of Het4 (Figure 1A, C).

Given the ability of Het4 to complement the weak CCM of PCC 73102, we were anticipating an N for C exchange between the diazotrophic cyanobacterium and Het4. Therefore, we first investigated protein categories representing carbon fixation, the carbon concentrating mechanism and nitrogen fixation. While the large and small subunit of RubisCO and phosphoribulokinase (Npun_F2752) were only slightly upregulated in the co-culture, we found a pronounced upregulation of three carbonic anhydrases (CAs) (Npun_F1420, Npun_R4176, Npun_F3687), one bicarbonate transporter (Npun_R2356), and one CO_2_ uptake transporter (Npun_F3690) (Figure 2). This observation supports the hypothesis that Het4 indeed provides CO_2_ to the cyanobacterium, especially because the strongest upregulation was observed for the predicted periplasmic CA (Npun_F3687) (Figure 2). In contrast, proteins involved in N-fixation behaved somewhat contrary to our assumptions. While the nitrogenase NifH itself was slightly upregulated in co-culture, accessory proteins such as the iron-molybdenum cofactor biosynthesis proteins NifN and NifE, and the stabilizing protein NifW, were clearly down-regulated. Even more surprising was the strongly reduced expression of heterocyst glycolipid biosynthesis proteins (Npun_R0038-R0043) pointing to a downregulation of heterocyst formation and N-fixation. The unexpected nature of these findings raises some questions on the nitrogen source under the given diazotrophic conditions. Notably, we observed an upregulation of two cyanophycinases (Npun_F1821 and Npun_R0196) suggesting that the potential impairment of nitrogen fixation is at least partially compensated by the utilization of nitrogen storage compounds (Figure 2).

**Figure 2.**
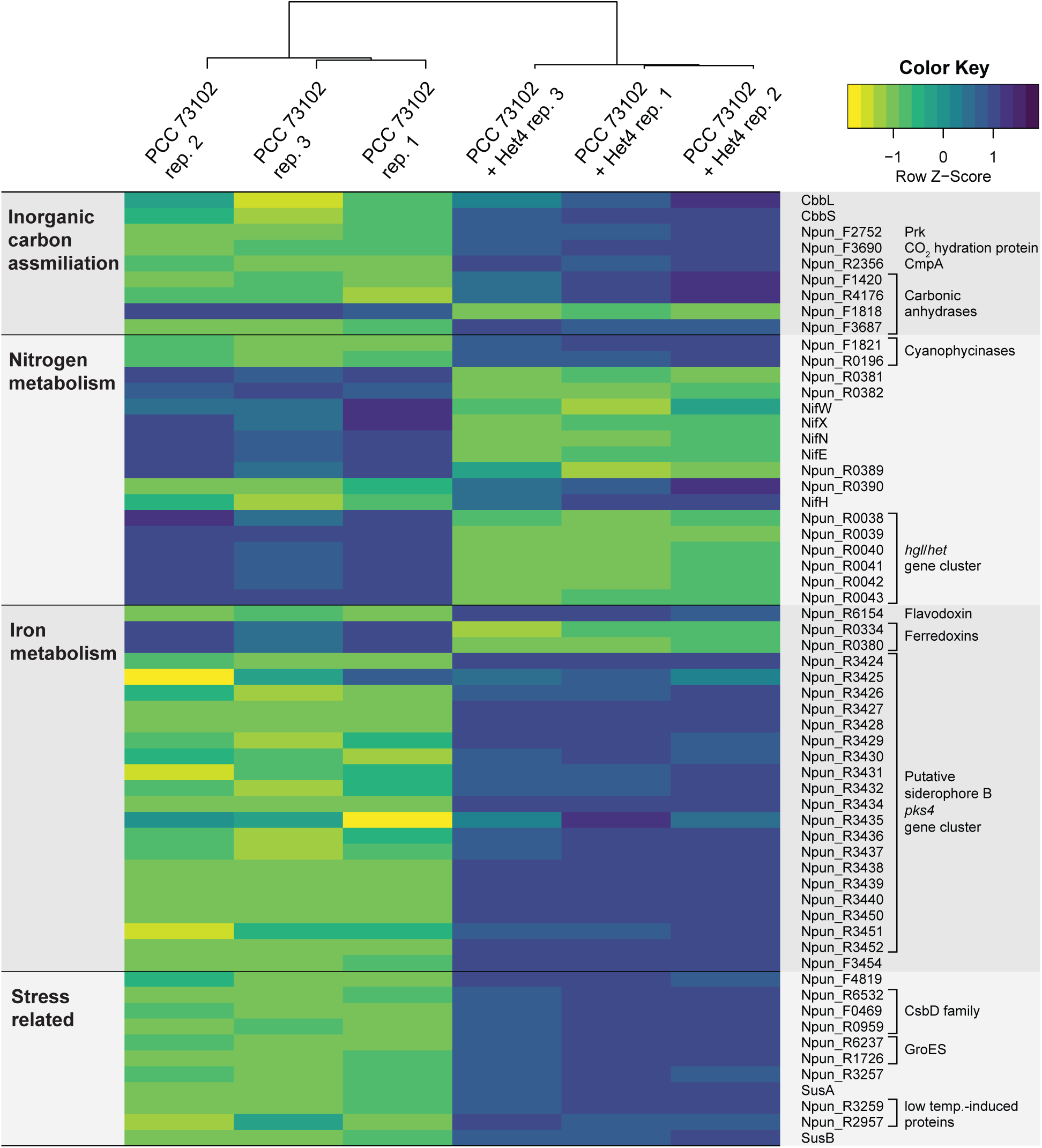
Heatmap illustrating the differential expression of selected proteins between *Nostoc punctiforme* PCC 73102 monoculture and co-culture with *A. tumefaciens* Het4. Rows represent individual proteins, while columns represent biological replicates of each condition. Proteins are labelled with gene names (right) and grouped by predicted cellular function (left). Log2 protein expression levels were transformed with a z-score normalization and represented with a color scale that varies from yellow (most downregulated) to blue (most upregulated) relative to the mean expression across all samples in each row. Hierarchical clustering was performed on samples and dendrograms indicate the similarity between them.

In the search for possible reasons for the partial downregulation of N-fixation, we were particularly struck by the upregulation of flavodoxin (Npun_R6154), which was the top hit among the upregulated proteins and is a marker for iron limitation (Sandmann et al., 1990) (Figure S5). Flavodoxins function analogously to ferredoxins as electron transfer proteins, but unlike ferredoxins are not dependent on iron. The parallel downregulation of two of the ferredoxins (Npun_R0334 and Npun_R0380) suggests a functional replacement of ferredoxins by flavodoxin in co-culture due to iron deficiency (Figure 2). This hypothesis was further fueled by the observed upregulation of the cryptic siderophore cluster PKS4 (Npun_R3414-Npun_R3453) (Liaimer et al., 2011) and its TonB-dependent siderophore receptor (Npun_R3454) (Figure 2B). These data point to a competition for iron between PCC 73102 and Het4. As adverse effects of iron-limitation on nitrogen fixation are well known (Larson et al., 2018), this competition may account for the preferential use of stored nitrogen over nitrogen fixation. The comparative proteomic study also revealed further evidence of negative interactions between the strains. For example, a number of stress markers were over-accumulating in the co-culture, in particular proteins of the cold shock protein family CsbD (Npun_R6532, Npun_F0469 and Npun_R0959), but also markers for osmotic stress such as a sucrose synthase (SusA and SusB) and chaperones of the GroES family (Npun_R6237 and Npun_R1726) (Figure 2). These findings only make it more astonishing that, overall, the dominant interaction is the mutualistic growth promotion of PCC 73102. Under standard BG11_0_ growth conditions, and without heterotrophic support, PCC 73102 shows severe signs of starvation evident by phycobiliprotein degradation. However, if the C-limitation is compensated by high carbonate, the negative effects of the co-cultivation dominate (Figure S3).

### Phenotypic response to inorganic carbon limitation in N. punctiforme PCC 73102 in mono- and co-cultures

Given that the strong dependence of the cyanobacterium on heterotrophs can primarily be explained by a weak CCM, we initiated phenotypic studies on the subcellular localization of RubisCO using immunofluorescence microscopy (IFM) and transmission electron microscopy (TEM). IFM with an antibody against the RubisCO subunit RbcL revealed a pronounced phenotypic variability in the subcellular localization of RbcL at all studied conditions. Three different localizations of the studied enzyme were observed: intracellular, membrane-near and extracellular RbcL (Figure 3). Overall, we found little evidence for carboxysomal localization of RubisCO. To exclude methodological bias, we also labelled the carboxysomes with a CcmK antibody and the small subunit of RubisCO with an RbcS antibody. While the typical carboxysomal structures were observed with the CcmK antibody, RbcS was mostly detected near the membrane thereby closely resembling the RbcL pattern (Figure S6). Even though RubisCO may be less accessible to the antibody in the carboxysomes, it can be clearly stated that large amounts of RubisCO are localized outside the carboxysome in PCC 73102 under the conditions tested. Control experiments with an anti-GFP antibody showed no signals at any of the above sites (Figure S7), confirming the specificity of the RbcL signal. In axenic cultures, RbcL was found mainly in the cytoplasm of the cells as well as near the cell membrane, but extracellular RbcL was also identified to some extent (Figure 3A and B, upper panel). In co-cultures with Het4, RbcL was more commonly detected in the extracellular sheath and, at lower amount, intracellularly (Figure 3A and B, lower panel). Interestingly, phenotypic plasticity of RbcL localization was also seen among filaments located next to each other, indicating strong heterogeneity within the culture under nitrogen deficiency (Figure 3A).

**Figure 3.**
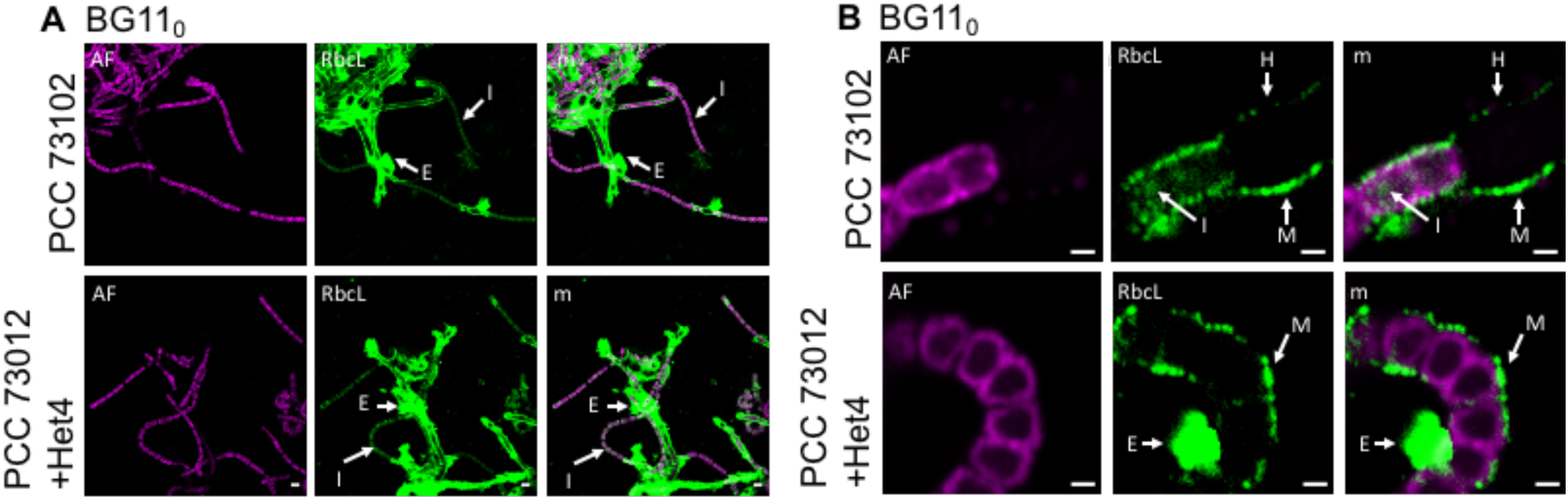
Immunofluorescence microscopy (IFM) images showing subcellular localization of RbcL, the large subunit of RubisCO, in *N. punctiforme* PCC 73012 monocultures and co-cultures with *A. tumefaciens* Het4 under nitrogen-deplete (BG11_0_) conditions. (A) shows overview micrographs illustrating the phenotypic heterogeneity among filaments either with predominant extracellular (E) or intracellular localization (I) of RubisCO. (B) depicts selected detail images showing subcellular localization of RbcL in mono- and co-cultures. Three localization types, extracellular, near membrane and cytosolic RbcL were found in both cultures with and without heterotrophic bacterium, however extracellular RbcL was more abundant in co-cultures, especially under nitrogen-deplete conditions. M = membrane-near RbcL, I = intracellular RbcL, E = extracellular RbcL H = heterocyst, AF = autofluorescence, m = merged. Scale bar (A): 5 µm; scale bar (B): 1 µm

To confirm the extracellular accumulation of RubisCO, we performed a second proteomic study in which we characterized the extracellular proteome of PCC 73102 using the recently described EXCRETE method (Russo et al., 2024). In total, 2970 proteins were identified in the PCC 73102 monoculture and 3085 were identified in co-culture with Het4. 2896 proteins were identified in both conditions. Overall, the large number of proteins identified in the exoproteome under BG11_0_ conditions, and the major overlap with the endoproteome, indicated a possible lysis of part of the PCC 73102 cells (Figure S8 and Dataset S2). Specifically, RubisCO was detected among the ten most abundant proteins in the medium, both in mono- and co-culture with Het4 along with further abundant intracellular proteins such as phycobilisome subunits and the periplasmic carbonic anhydrase (Fig. S8). The strong extracellular accumulation of RubisCO is apparently specific for N- and C-limiting conditions, since a recently published exoproteomic study of PCC 73102 showed that under N- and C-replete conditions predicted extracellular proteins were enriched, but not RubisCO (Russo et al., 2024). To evaluate whether the exoproteomic difference between nitrogen deplete (BG11_0_) and nitrogen replete (BG11) conditions is also reflected in the extracellular localization of RubisCO detected by IFM, we also transferred mono- and co-cultures from BG11_0_ conditions to BG11 conditions and evaluated them by IFM after eleven days (Figure S9). Indeed, the apparent proportion of external RbcL decreased, while the intracellular RbcL content in vegetative cells increased thereby supporting the findings of the exoproteomic analyses (Figure S9 and Russo et al., 2024). This phenomenon suggests that RubisCO is primarily accumulating outside the cells under inorganic carbon and nitrogen deficiency.

Notably, RubisCO was not differentially accumulating in the exoproteome of the mono- and co-culture with Het4 (Figure S8 and Dataset S2). We assume that part of the extracellular RubisCO which sticks to the cell-bound mucus layer especially in the co-culture (Figure 3A) is not enriched by the exoproteomic method used here but remains in the cellular pool. Nonetheless, the characterization of the exoproteome provides further evidence that *N. punctiforme* suffers from starvation under standard BG11_0_ growth conditions and may sacrifice cells and proteins including RubisCO to increase the viability of the community.

To gain further insight into ultrastructural differences within cyanobacterial cultures and between axenic and co-cultures under inorganic carbon limitation, we used TEM. Interestingly, there was structural heterogeneity with respect to the capsular sheath (CPS) among PCC 73102 filaments within and between axenic and co-cultures (Figure 4). The presence of the capsular sheath varied, with some filaments showing no detectable sheath, while others had a relatively electron-lucent or compacted sheath that was up to 2 µm thick. We had the impression that the capsular sheath of PCC 73102 was more prominent in co-cultures. In addition, we observed small electron-dense granular particles outside the PCC 73102 cells, often associated with the outer surface of the capsular sheath, and these particles were present at far larger amount in co-cultures (Figure 4B). Sheathed cells contained an increased number of small electron-dense granular particles outside the cells. The presence of external RbcL in the granular particles was confirmed by pre-embedding immunogold TEM (Figure 5). In co-cultures, large amounts of extracellular RbcL could be detected in the periphery of the cyanobacterial sheath, similar to the IFM studies under the same conditions (Figure 3 and 5). Control experiments with an anti-GFP antibody showed no signals at any of the above sites (Figure. S10)

**Figure 4.**
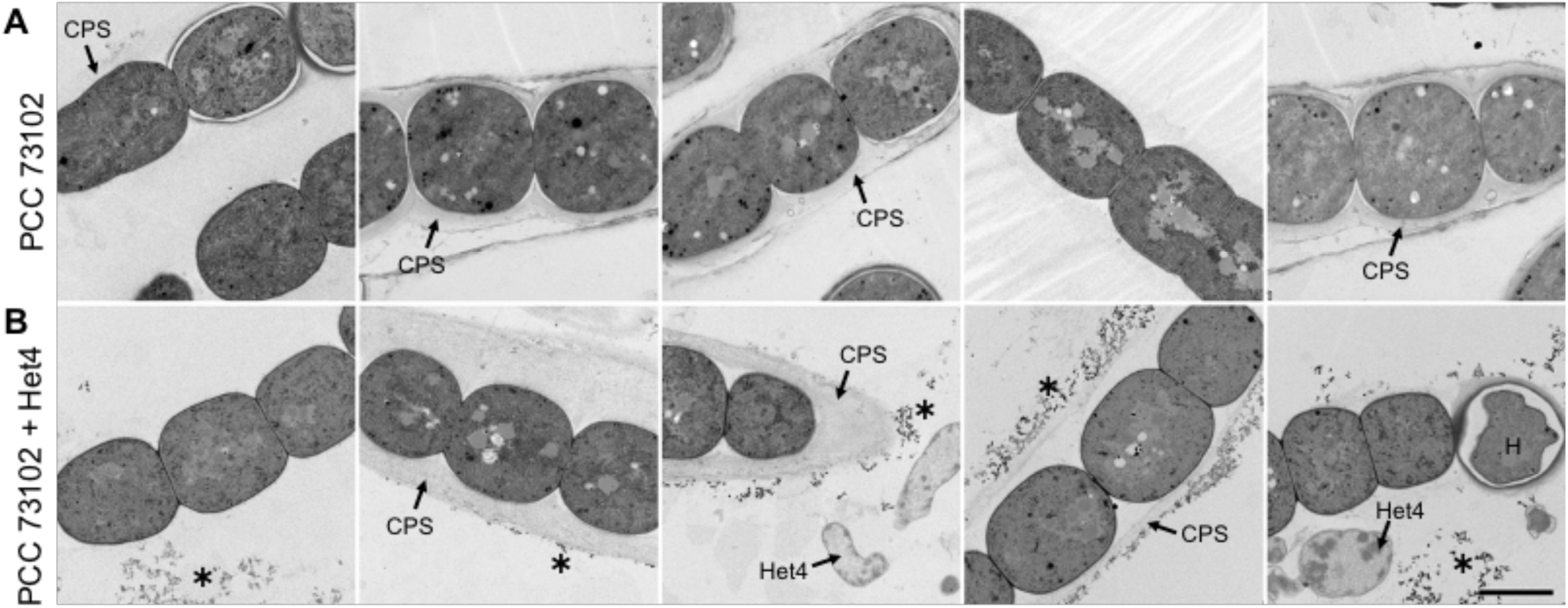
Transmission electron microscopy (TEM) images of *N. punctiforme* PCC 73102 grown in (A) monoculture or (B) co-culture with *A. tumefaciens* Het4 under nitrogen-deplete conditions. (A) TEM images of axenic *N. punctiforme* PCC 73102 show heterogeneity in the polysaccharide capsule. (B) TEM images of *N. punctiforme*-Het4 co-cultures show heterogeneity in the polysaccharide capsule and an accumulation of granular material (highlighted by an asterisk) predominantly in the periphery of the polysaccharide layer. CPS = capsular polysaccharides, H = heterocyst. Scale bar = 2 µm

**Figure 5.**
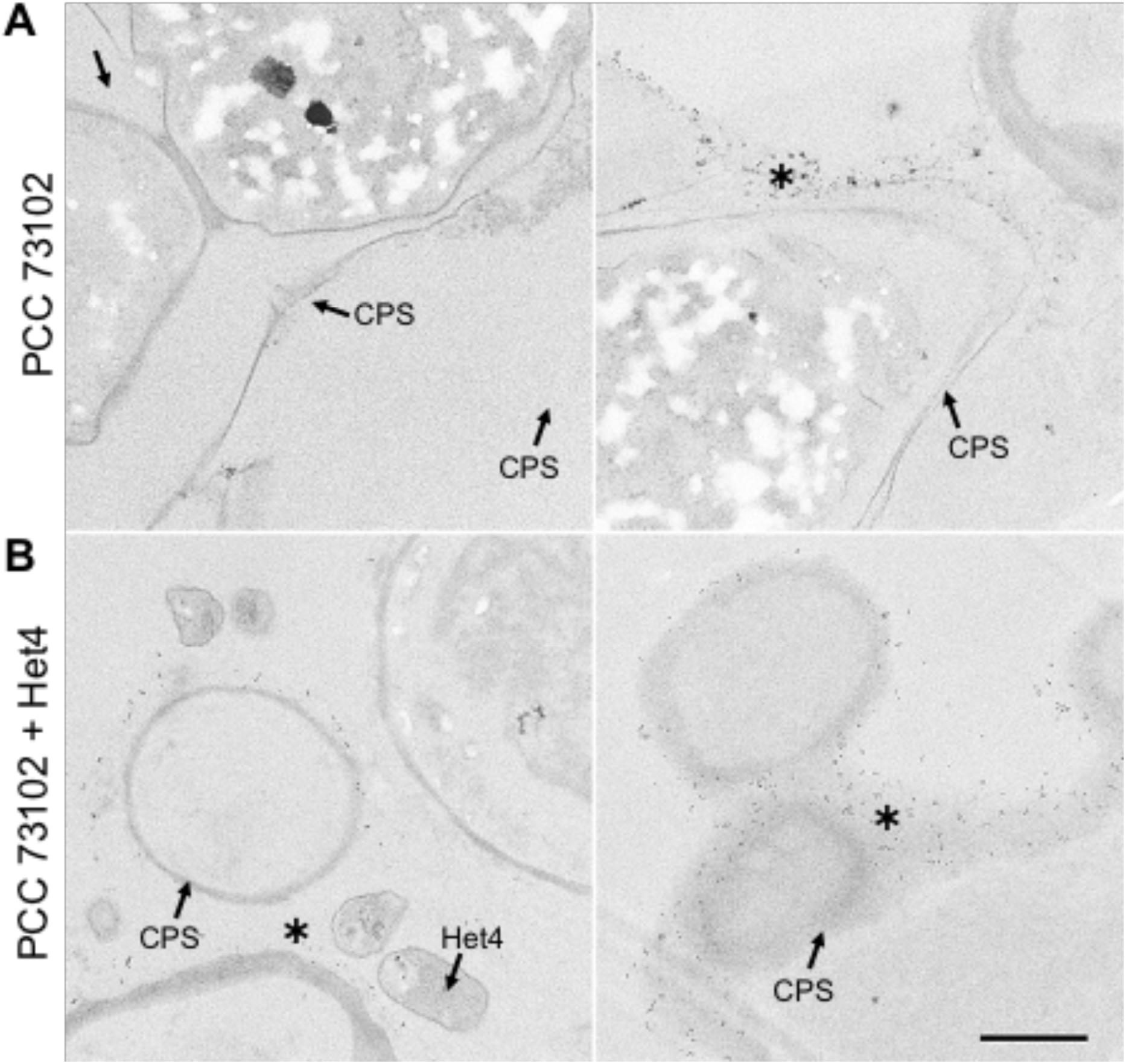
Immunogold TEM images of (A) *N. punctiforme* PCC 73102 monoculture and (B) PCC 73102-Het4 co-culture samples incubated with antibody against RubisCO subunit RbcL. Immunogold particles are visible at the periphery of the polysaccharide capsule (CPS) in both mono- and co-culture (indicated by an asterisk), but they are much more abundant in the co-culture with Het4. Scale bar: 1µm

## Discussion

The present study not only provides detailed insights into the intimate relationship of the symbiotic strain *N. punctiforme* PCC 73102 and its natural heterotrophic partner *Agrobacterium tumefaciens* Het4, but also demonstrates the limited autonomy of the diazotrophic photoautotrophic cyanobacterium under N- and C-deficient conditions. We showed that a combined C and N limitation enforces an almost obligate dependence of PCC 73102 on heterotrophic partners. It is particularly remarkable that C limitation manifests itself already at the commonly used inorganic carbon concentrations of the BG11_0_ standard medium. Under these conditions, the axenic strain PCC 73102 shows clear signs of bleaching and a pronounced accumulation of intracellular proteins in the exoproteome, that likely results from starvation and cell lysis.

The fact that the dependence on heterotrophic bacteria can be abolished by addition of high carbonate concentrations (Figure 1A) indicates a weak CCM in *N. punctiforme* compared to model cyanobacteria such as *Synechocystis* sp. PCC 6803. In the search for causes, we discovered differences in the genetic repertoire of bicarbonate uptake transporters, in particular a lack of the gene encoding the high-affinity uptake transporter SbtA. Furthermore, we detected a predominantly extracarboxysomal localization of RubisCO, often in the cytoplasmic membrane region. Both findings are similar to what is known for the bloom-forming freshwater cyanobacterium *Microcystis aeruginosa* PCC 7806 (Barchewitz et al., 2019). *N. punctiforme* and *Microcystis* are among the many cyanobacteria that are very difficult to axenize and maintain as axenic strains (Heck et al., 2016; Alvarenga et al., 2017; Warshan et al., 2018a). Notably, the Pasteur Culture Collection recommends the addition of NaHCO_3_ to standard BG11/BG11_0_ medium for both *Microcystis* and *N. punctiforme* for axenic maintenance (https://catalogue-crbip.pasteur.fr/). It is tempting to speculate that these cyanobacteria are more specialized than unicellular model cyanobacteria on utilizing respiratory CO_2_ from heterotrophic bacteria and are therefore less dependent on a strong CCM in nature. The cytoplasmic membrane localization of RubisCO may facilitate the intimate inorganic carbon supply. This hypothesis is supported by the proteomic findings in the present study. In particular, the activation of carbonic anhydrases and bicarbonate or CO_2_ transporters in co-cultures with Het4 is striking. The benefits of this inorganic carbon feeding by Het4 are clearly evident through growth promotion, even against the background of negative interactions, including competition for iron (Figure 6).

**Figure 6.**
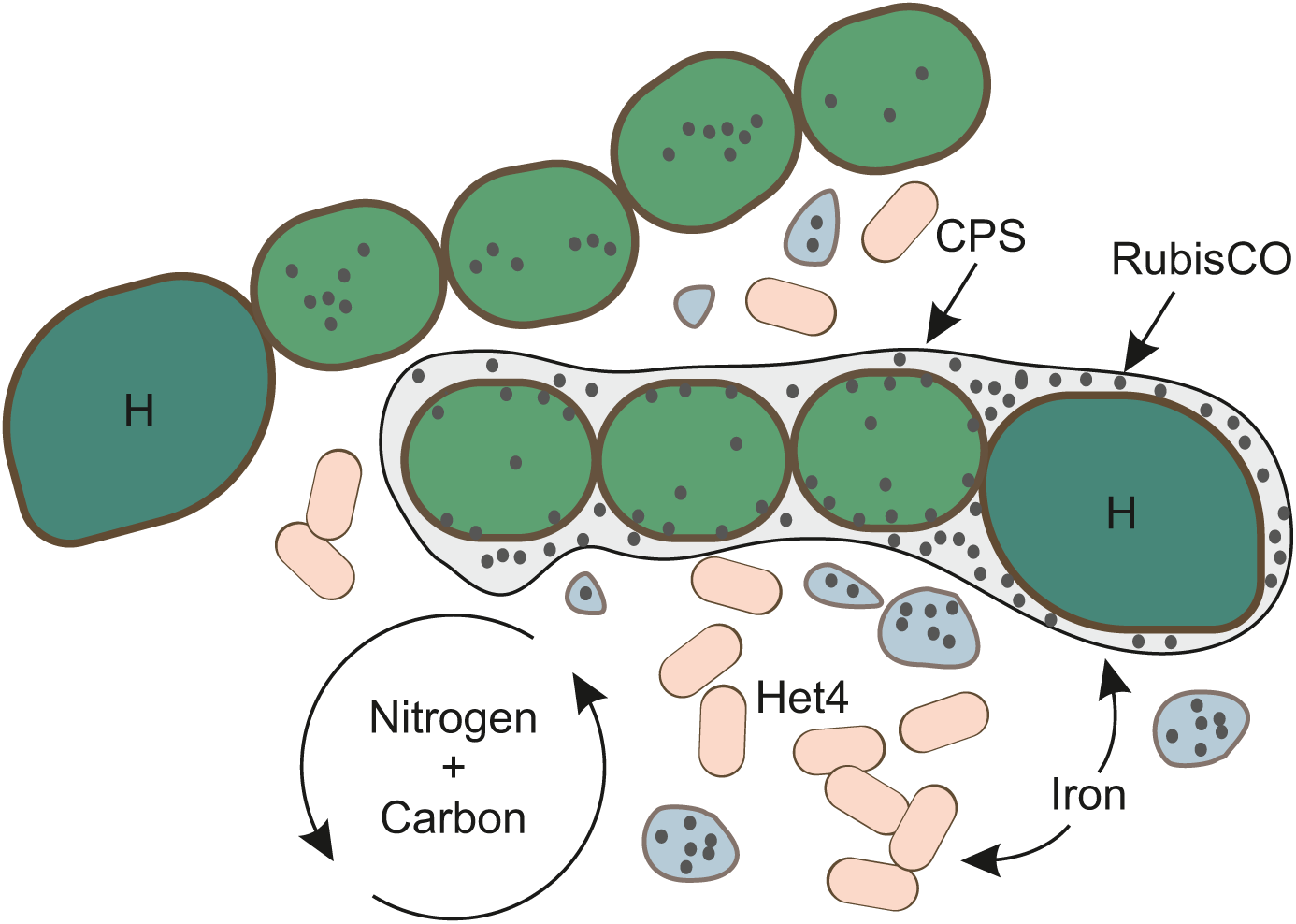
Model for the interaction between *N. punctiforme* PCC 73102 and *A. tumefaciens* Het4 under N- and C limiting conditions.

In addition to its dependence on inorganic carbon supply by heterotrophic partners, a pronounced phenotypic plasticity of the PCC 73102 filaments is also noteworthy in the present study. This plasticity was observed with regard to the extent and nature of the extracellular sheath as well as with regard to the subcellular localization of RubisCO and the accumulation of RubisCO-containing extracellular granules. The extracellular sheath of microalgae is commonly referred to as the phycosphere and provides a nutrient-rich microscale interface favoring phototroph-heterotroph interactions (Seymour et al., 2017). The plasticity may reflect a division of the population into filaments specializing on cooperation with Het4 and those that thrive independently of it. Division of labor and cheating on resources are common phenomena in mono- and multispecies biofilms and may also contribute to versatility and resilience of the *Nostoc* population (van Gestel et al., 2015; Martin et al., 2020). As we have no global proteomic insight into the differences between the distinct filament types, we can only speculate about traits favoring cooperation. The increased accumulation of RubisCO-containing extracellular granules around sheathed filaments in the co-culture may indicate a major role of protein recycling for the cooperation between PCC 73102 and Het4. The denitrification potential of Het4 may contribute to the N recycling process. Notably, an enrichment of denitrifying heterotrophic bacteria was also observed in diazotrophic freshwater cyanobacteria of the genus *Dolichospermum* under N-limitation (Pascault et al., 2021).

The extracellular mucus layer may also be one of the reasons for the specificity of *Nostoc* interactions with its heterotrophic microbiome. Even though we have studied only two field strains, the observed major differences in their heterotrophic communities support a host specificity in principle. Previous studies have suggested that Rhizobiales generally play important roles in plant-cyanobacterium consortia, and our data strongly support these findings (de Vries and de Vries, 2022). Yet, the impact of *Rhizobium*/*Agrobacterium* on PCC 73102 was clearly strain-specific as the closely related Het6 isolate did not show signs of growth promotion (Figure S2). The bilateral selectivity of the growth promotion indicates that variable functional traits, that are not part of the species’ core genome, may be crucial for the interaction.

## Conclusion

The present study sheds new light on the dependence of symbiotic cyanobacteria of the genus *Nostoc* on respiratory activities of associated heterotrophic bacteria. This dependence is likely enforced by a weak CCM and is promoted by a physical interaction around the pronounced sheath of the cyanobacteria. Symbiotic *Nostoc* strains apparently rely on partners for the supply of carbon, either heterotrophic bacteria in free-living *Nostoc* colonies that provide CO_2_ or plant hosts that provide organic carbon. Although the mutualistic interactions of *Nostoc* and *Agrobacterium* have been observed mainly under N and C deficiency, it can be assumed that they play a major role in many habitats and strongly influence the vitality of *Nostoc* colonies. Understanding the interdependencies is not only important for assessing the role of phototroph-heterotroph interactions in nutrient cycles, but also central for the biotechnological exploitation of *Nostoc* strains.

## Materials and methods

### Strains and growth conditions

Axenic *Nostoc punctiforme* PCC 73102 (hereafter PCC 73102) was obtained from the Pasteur Culture Collection and maintained in liquid BG11_0_ medium under continuous illumination of 25 µmol photons m^-2^s^-1^ at 23°C. *Nostoc* sp. KVJ2 (hereafter KVJ2) and *Nostoc* sp. KVJ3 (hereafter KVJ3) were isolated from Northern Norway as described previously (Liaimer et al., 2016) and since then cultivated diazotrophically in BG11_0_ medium under continuous light of 25 µmol photons m^-2^s^-1^ at 23°C. Heterotrophic bacteria (Het1-6) were isolated from the strains KVJ2 and KVJ3 by plating the culture on R2A agar plates and purified over several cultivation cycles at 23°C.

Effects of inorganic carbon concentrations on the growth of cyanobacteria were studied in three different carbonate concentrations: low (BG11_0_ without added Na_2_CO_3_), medium (BG11_0_) and high (BG11_0_ enriched with 10x higher Na_2_CO_3_ than in standard BG11_0_). Concentration of the sodium ions in low bicarbonate medium was adjusted by NaCl. Prior to the experiment, PCC 73102, KVJ2 and KVJ3 cyanobacteria were washed twice using low carbonate BG11_0_ medium, resuspended in same medium (OD750=4.4) and a volume of 10 µL each was dropped onto three agar plates in three biological replicates. To study the growth promotion of heterotrophic bacteria, heterotrophic bacteria were washed and resuspended in a similar concentration as cyanobacteria (final OD600 = 2.2) and 10 µL of the bacteria were added on the top of the cyanobacterial drop (physical interaction) at 23°C.

### Cultivation in liquid cultures

Prior to the experiment, PCC 73102 was washed twice using BG11_0_ medium and resuspended in 20 mL BG11_0._ The washed cells were split into two flasks, each containing 10 mL cell suspension. In parallel, a single colony of Het4 was picked from a R2A (CL01.1, Carl Roth, Karlsruhe, Germany) plate, transferred to 4 mL LB medium (X968.4, Carl Roth, Karlsruhe, Germany) and incubated at 28 °C, 210 rpm shaking for 14 h. The cells were harvested, washed twice with BG11_0_ medium, resuspended in 2 mL BG11_0_ medium (OD600 = 2.3) and added to one of the PCC 73102 cultures.

Both cultures were then kept under continuous illumination of 40 µE without shaking for 7 days at 23°C. Samples for immunofluorescence microscopy were taken on day 1, day 4 and day 7 by aseptically removing 1 mL cell suspension and replenishing to the original volume with fresh medium. Cells were fixed as described below and stored at -20°C in PBS (140 mM NaCl, 2.7 mM KCl, 8.0 mM Na_2_HPO_4_, 1.8 mM KH_2_PO_4_, pH = 7.5) with 10% w/v glycerol.

On day 7, 5 mL of each culture was washed twice with BG11 medium and resuspended in 10 mL BG11 medium while replenishing the lost volume with BG11_0_ medium. All four cultures were then kept under the same conditions as before for another 7 days. Again, samples were taken, fixed and stored on day 8, day 11 and day 14 as described above.

### DNA sequencing

For Sanger sequencing of the heterotrophic bacterial isolates and for Illumina MiSeq 16S rRNA gene amplicon sequencing, the total DNA from KVJ3 and KVJ2 cultures were isolated using a DNA Isolation Kit for Cells and Tissues (Sigma-Aldrich, Taufkirchen, Germany). Universal primers 27F and 1492R were used in the amplification and Sanger sequencing of single bacterial isolates at LGC Genomics GmbH, Berlin, Germany. V3 and V4 regions of the 16S rRNA gene were PCR amplified and 16S rRNA gene amplicons were sequenced by Illumina MiSeq platform at the Institiute of Biology, University of Helsinki, Finland. High-quality reads between 10-471 bp with unambiguities were analyzed using Mothur v1.39.5 (Schloss et al., 2009) and aligned against the Silva database (Release 138) (Quast et al., 2013) with kmer size of 8. Maximum-likelihood phylogenetic tree with bootstrapping of 100 iterations was constructed using the MEGA X software (Tamura et al., 2021). For the whole genome PacBio and Illumina hybrid sequencing, high-molecular weight DNA was isolated from KVJ3 and *Agrobacterium tumefaciens* Het4 using phenol-chloroform extraction. Cells were broken using lysozyme in buffer containing 50 mM Tris-HCl, 100 mM EDTA, 0.1 M NaCl (pH=7) at 37 °C for one hour. Proteinase K and RNAase were added on the cells and incubation was continued at 56 °C for 2 hours. DNA was extracted and purified with phenol and chloroform and precipitated with 70% ethanol in the presence of 0.1 M of NaCl. DNA was dissolved in TE buffer and sequenced at the Institute of Biology, University of Helsinki, Finland. PacBio RSII reads were assembled using HGAP3 implemented in SMRT portal with a default genome size of 6 Mbp and circulated using Gap4. Illumina MiSeq reads were filtered by cutadapt v 1.14, m=100, q=25 (Martin, 2011) and genome was polished using bwa v0.7.12-r1039 (Li and Durbin, 2009) and Pilon v1.16 (Walker et al., 2014) with default parameters. Newly sequenced genomes were annotated using NCBI prokaryotic genome annotation pipeline (PGAP) (Tatusova et al., 2016).

High-quality reads were mapped using BWA-MEM (v0.7.17.1) (Li and Durbin, 2009). Reads <20 bp, secondary aligned reads and PCR duplicates were removed using BAM filter (v0.5.9) (Li et al., 2009) and features were called using featureCounts (v1.6.4) (Liao et al., 2014) with paired-end mode and exclusion of chimeric structures. Reads mapped to Het4 genome were removed from the differential gene expression analysis. DESeq2 (v1.28.1) (Love et al., 2014) with rlog transformation was used to call the differentially expressed (DE) genes.

### Preparation of endoproteome and exoproteome fractions

To obtain the exoproteome fraction, cultures were centrifuged for 10 min at 5000 × g. The supernatant was removed, centrifuged again for 10 min at 10 000 × g and transferred to a fresh microcentrifuge tube. Pre-cultures for the preparation of endoproteomics co-cultivation plates were grown and washed as described above. Subsequently, three 50 µL droplets of either PCC 73102 were spotted on freshly prepared BG110 plates. For the cocultivation plates, 50 µL of Het4 were gently dropped onto the PCC 73102 spots. Plates were incubated for 16d under the same conditions as described above. Endoproteome fractions were obtained by scraping culture spots from their respective agar plates and then resuspending the biomass in 300 µL of lysis buffer (25 mM Tris-HCl, 5% (w/v) glycerol, 1% (v/v) Triton X-100, 1% (w/v) sodium deoxycholate, 0.1% (w/v) sodium dodecyl sulfate (SDS) and 1 mM EDTA). Zirconium oxide beads (diameter 0.15 mm) were added to the cell suspension and cells were broken in a Bullet Blender Storm 24 (Next Advance) with three cycles of 5 min. Cell lysates were centrifuged at 10 000 × g for 10 min and the resulting supernatant transferred for further analysis. The protein content of all samples was determined with a PierceTM BCA assay kit (Thermo Fisher Scientific).

### Proteomic analysis

Sample preparation for proteomic analysis was done according to the EXCRETE workflow (Russo *et al*., 2024). Briefly, the equivalent of 10 µg of protein was harvested from the supernatant and transferred to 2 mL microcentrifuge tubes. NaCl and SDS were added to a final concentration of 10 mM and 1% (w/v) and samples were reduced with 5 mM TCEP and alkylated with 5.5 mM CAA. Protein aggregation was induced by adding LC-MS grade ethanol to a final concentration of 50% (v/v) followed by SiMAG-Carboxyl magnetic particles (product no. 1201, Chemicell) to a final concentration of 0.5 µg µL^-1^. Samples were incubated for 10 min with shaking at 1000 rpm and, subsequently, magnetic particles were separated on a magnetic rack and washed, on-magnet, 3 times with 80% (v/v) ethanol. Protein digestion was done overnight at 37°C on-bead in 100 µL of 25 mM ammonium bicarbonate containing 0.5 µg of MS grade Trypsin/LysC (Promega) (enzyme/protein ratio of 1:20 (w/w)). Following protein digestion, magnetic particles were separated for 60 s and supernatants were recovered. Peptide purification and desalting, LC-MS analysis and raw data processing were done as previously described (Russo *et al*., 2024). For protein identification a protein group was considered identified when it was present in at least 70% of the replicates with a minimum of three replicates. Missing values were imputed using the k-nearest neighbors’ algorithm. Proteins were annotated using EggNOG v5.0 (Huerta-Cepas *et al*., 2019), PsortB v3.0 (Yu *et al*., 2010), SignalP 6.0 (Teufel *et al*., 2022) and UniProtKB (The UniProt Consortium, 2023). Data analysis and visualization were performed using custom scripts in R (4.3.0) with packages ggrepel (0.9.5), ggplot2 (3.5.1), viridis (0.6.5), readxl (1.4.3) and gplots (3.1.3.1). Differential protein analysis was done using a Student’s two-sample unpaired t-test with permutation-based multiple test correction with a cutoff criterion of fold change = 2 and adjusted p-value < 0.05.

### Immunofluorescence microscopy

Cells were harvested by centrifugation and washed twice in freshly prepared PBS. Washed cells were resuspended in 4% w/v paraformaldehyde in PBS (J61899.AK, Thermo Fisher, Hennigsdorf, Germany), incubated for 10 min at RT, washed two more times with PBS, resuspended in ultra-pure water and subsequently spread on 2H coverslips (01-0012/2, Langenbrinck, Emmendingen, Germany) to air-dry until sufficient attachment was achieved without completely desiccating the cells. All following steps were carried out in a humidifier.

The specimens were washed once with PBS for 5 min, then permeabilized using 2 mg/mL of lysozyme (0663-10G, VWR, Darmstadt, Germany) in PBS with 0.3% w/v Triton-X 100 (PBS-TX) for 30 min at RT, washed twice in PBS-TX for 3 min, blocked using 1% w/v Polyvinylpyrrolidone K30 (4607.1, Carl Roth, Karlsruhe, Germany) in PBS with 0.3% w/v Tween-20 (PBS-T) for 1h at 4°C and washed twice in PBS-T for 3 min.

For antibody labeling, the specimen was incubated for 1h at RT with primary antibodies (rabbit anti-rbcL large subunit, form I polyclonal antibody, AS03 037A, Agrisera, Vännas, Sweden; for negative controls: rabbit anti-GFP N-terminal polyclonal antibody, Sigma-Aldrich, G1544) diluted to 4 µg/ml in PBS-T, and washed twice in PBS-T for 3 min.

Secondary antibodies (goat anti-rabbit IgG Alexa Fluor 488, A-11008, Thermo Fisher, Hennigsdorf, Germany) were diluted 1:200 in PBS-T and applied by incubation for 1h at 4°C in the dark. After two final washes in PBS-T, specimens were air-dried and mounted on slides using ProLong Glass (P36980, Thermo Fisher, Hennigsdorf, Germany). Slides were imaged using a Zeiss LSM780 (Carl Zeiss Microscopy, Jena, Germany) laser scanning confocal microscope equipped with a Plan-Apochromat 63x/1.40 oil immersion lens. Alexa Fluor 488 was excited at 488 nm and detected at 493 to 550 nm, whereas autofluorescence was excited at 633 nm, and detected between 647 and 687 nm.

### Electron microscopy

Cells from 2 mL of PCC 73102 culture were collected, washed, and fixed by 2.5% glutaraldehyde, 2.0% formaldehyde in 0.1 M Na–cacodylate buffer, pH 7.4 similarly as described earlier (Barchewitz et al., 2019). Subsequently, samples were overlaid by a thin layer of 1 % low-melting agarose, dehydrated in a graded EtOH series and acetone and embedded in low viscosity resin (Agar Scientific, Stansted, Essex, UK). Ultrathin sections stained with uranyl acetate and lead citrate were examined in a Talos F200C transmission electron microscope (Thermo Fisher Scientific, Waltham, MA, USA), operated at 200 kV.

### Immunogold Electron Microscopy

Cells were carefully removed from agar plates, resuspended in 1 mL PBS, and treated as described previously in section “Immunofluorescence microscopy” until secondary antibody hybridization. After washing in PBS, specimens were incubated for 60 min with 10-nm-gold-conjugated goat-anti-rabbit IgG (Sigma-Aldrich, product number G7402), diluted 1:25 in PBS, and washed in PBS for 2 x 5 min. After a further 5-min-washing step in PBS supplemented with 0.5 M NaCl, specimens were incubated for 5 min in 0.1 M Na-phosphate buffer (pH 7.3), post-fixed with phosphate-buffered 3% glutaraldehyde for 30 min, washed in phosphate buffer for 5 min, and dehydrated in a graded EtOH series and acetone. Coverslips were then embedded in Spurr low viscosity resin (Science Services, Munich, Germany) as described in detail previously (Batsios et al., 2013). After polymerization, coverslips were removed from the polymerized resin by several cycles of cooling (liquid N_2_) and re-warming to room temperature. Areas with bacteria were selected by phase contrast microscopy, mounted (Batsios et al., 2013), sectioned on an ultramicrotome at a thickness of 90 nm, stained with uranyl acetate for 5 min, and analyzed in a Talos F200C.

## Acknowledgements

The study was supported by Academy of Finland grant No. 332215 to JET, DFG Project No. 406260942 (cryo-TEM) to Petra Wendler and DFG Project No. 239748522-SFB 1127 (DAR, JAZZ and ED). We acknowledge the DNA Sequencing and Genomics Laboratory, Institute of Biotechnology, University of Helsinki for sequencing. and Florian Meier and Denys Oliinyk, Jena University Hospital, for assistance with the proteomic analysis.

## Data availability statement

The authors confirm that sequencing data supporting the findings of the study are publicly available in the NCBI database [SRA accession numbers SRR24556967 and SRR24556966 and project numbers PJNA599284 and PRJNA599316]. The mass spectrometry proteomics data have been deposited to the ProteomeXchange Consortium via the PRIDE partner repository with the dataset identifier PXD052511.

## Funding statement

The study was supported by Academy of Finland grant No. 332215 to JT, DFG Project No. 406260942 (cryo-TEM) to Petra Wendler and DFG Project No. 239748522-SFB 1127 to DAR, JAZZ and ED.

## Conflict of interest disclosure

None of the authors has declared a conflict of interest. Ethics approval statement: There are no ethical issues related to our manuscript.

## CRediT

ED and JET designed the work, JET, DAR, MH, OB conducted experiments, JET, DAR, MH and JAZZ analyzed data, AL provided resources, all co-authors contributed to the discussion and writing of the manuscript.

## Notes

### Competing Interest Statement

The authors have declared no competing interest.

