## Supplementary Information for "Competition and interdependence define multifaceted interactions of symbiotic *Nostoc* sp. and *Agrobacterium* sp. under inorganic carbon limitation"

**Table of contents:**

**Supplementary tables:**

**Table S1** Genome analysis of *N. punctiforme* PCC 73102 and *Nostoc* sp. KVJ2 and KVJ3 to identify genes related to the cyanobacterial carbon concentrating mechanism (CCM).
**Table S2** Results of the pairwise t-Tests conducted to determine significant differences in Surface Greenness (Figure S3).

**Supplementary figures:**

**Figure S1** Phylogenetic tree based on 16S rRNA V3-V4 regions.
**Figure S2** Heterotrophic bacterial isolates promote growth of *N. punctiforme* PCC 73102.
**Figure S3** Digital image analysis of growth promotion assays.
**Figure S4** Amino acid identity (AAI) of *A. tumefaciens* Het4 illustrated by AAI profiler.
**Figure S5** Volcano plot illustrating differential expression of all proteins identified in the *N. punctiforme* PCC 73102 endoproteome in monoculture and co-culture with *A. tumefaciens* Het4.
**Figure S6** Immunofluorescence labelling of carboxysomal shell protein CcmK and the small subunit of RubisCO RbcS in *N. punctiforme* PCC 73102.
**Figure S7** Negative control for immunofluorescence staining.
**Figure S8** Intensity of proteins identified in the *N. punctiforme* PCC 73102 exoproteome ordered by rank.
**Figure S9** Immunofluorescence detection of RbcL under nitrogen replete conditions in BG11 medium.
**Figure S10** Immunogold TEM. Co-culture of *N. punctiforme* PCC 73102 and *A. tumefaciens* Het4 was incubated with anti-RbcL and, for control, anti-GFP.

**Supplementary tables**

**Table S1** Genome analysis of *N. punctiforme* PCC 73102 and *Nostoc* sp. KVJ2 and KVJ3 to identify genes related to the cyanobacterial carbon concentrating mechanism (CCM).

| Gene | PCC 73102 | KVJ2 | KVJ3 | Reference |
|---|---|---|---|---|
| bicA | + | + | + | (Sandrini *et al*, 2014) |
| sbtA | - | - | - | (Sandrini *et al.*, 2014) |
| sbtB | - | - | - | (Sandrini *et al.*, 2014) |
| ccmR2 | + | + | + | (Sandrini *et al.*, 2014) |
| nhaS3 | + | + | + | (Sandrini *et al.*, 2014) |
| CmpA | + | + | + | (Sandrini *et al.*, 2014) |
| CmpB | + | + | + | (Sandrini *et al.*, 2014) |
| CmpC | + | + | + | (Sandrini *et al.*, 2014) |
| CmpD | + | + | + | (Sandrini *et al.*, 2014) |
| EcaA | - | - | - | (Sandrini *et al.*, 2014) |
| EcaB | - | - | - | (Sandrini *et al.*, 2014) |
| CcaA (icfA) | + | + | + | (So & Espie, 2005) |
| CcmM | + | + | + | (So & Espie, 2005) |
| NdhD3 | + | + | + | (Shibata *et al*, 2001) |
| NdhF3 | + | + | + | (Shibata *et al.*, 2001) |
| CupA | + | + | + | (Shibata *et al.*, 2001) |
| NdhD4 | + | + | + | (Shibata *et al.*, 2001) |
| NdhF4 | + | + | + | (Shibata *et al.*, 2001) |
| CupB | + | + | + | (Shibata *et al.*, 2001) |

**Table S2** Results of the pairwise t-Tests conducted to determine significant differences in Surface Greenness (Figure S3).

| Day | Condition | Δ(Mean Greenness) | Mean Greenness Nostoc | Mean Grenness Nostoc + Het4 | Test statistic | p Value | Degrees of Freedom |
|---|---|---|---|---|---|---|---|
| 0 | High | 2.52456 | 11.56156 | 9.03700 | 0.54690 | 5.93E-01 | 13.71682 |
| | Low | -12.18278 | 10.44856 | 22.63133 | -1.50679 | 1.65E-01 | 9.20178 |
| | Medium | 8.88878 | 19.89278 | 11.00400 | 1.12142 | 2.86E-01 | 11.23541 |
| 1 | High | 5.18933 | 11.54200 | 6.35267 | 1.87748 | 8.37E-02 | 12.65808 |
| | Low | -0.76300 | 1.71644 | 2.47944 | -0.98809 | 3.39E-01 | 15.25803 |
| | Medium | -21.49478 | 1.68133 | 23.17611 | -10.42848 | 4.10E-06 | 8.45300 |
| 3 | High | 50.90411 | 75.13944 | 24.23533 | 7.74294 | 5.65E-06 | 11.85538 |
| | Low | 24.70300 | 32.14044 | 7.43744 | 9.15936 | 9.76E-07 | 11.90168 |
| | Medium | 20.34489 | 25.14811 | 4.80322 | 11.35697 | 9.29E-08 | 11.94898 |
| 5 | High | 76.04422 | 124.90444 | 48.86022 | 10.63478 | 1.01E-07 | 12.80782 |
| | Low | 38.88156 | 43.91700 | 5.03544 | 10.84684 | 4.69E-07 | 10.55758 |
| | Medium | -18.67167 | 43.17433 | 61.84600 | -4.25610 | 6.04E-04 | 15.99711 |
| 7 | High | 47.31600 | 157.98589 | 110.66989 | 4.39186 | 4.90E-04 | 15.47384 |
| | Low | 31.18111 | 48.27478 | 17.09367 | 6.22785 | 7.73E-05 | 10.55796 |
| | Medium | -47.34278 | 57.12156 | 104.46433 | -7.04424 | 3.04E-06 | 15.73557 |
| 9 | High | 24.33922 | 179.83044 | 155.49122 | 3.50506 | 2.96E-03 | 15.89005 |
| | Low | -18.46256 | 37.53856 | 56.00111 | -3.35114 | 5.79E-03 | 11.96211 |
| | Medium | -86.88833 | 60.23356 | 147.12189 | -12.64963 | 3.24E-08 | 11.79074 |

### Supplementary figures

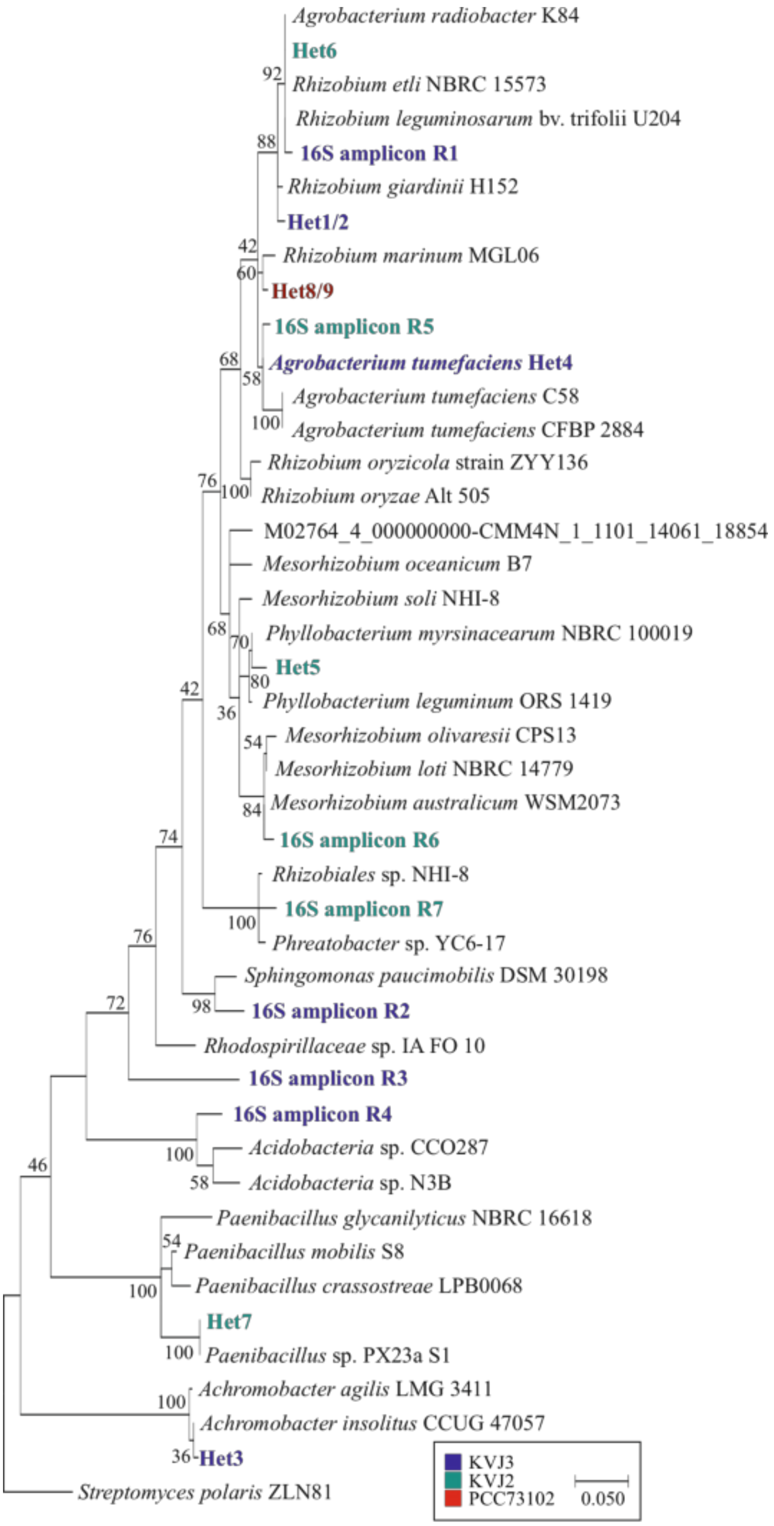


**Figure S1** Phylogenetic tree based on 16S rRNA V3-V4 regions. Most of the isolated heterotrophic bacteria belonged to the *Rhizobium*/*Agrobacterium* group. Het8/9 represent strains isolated from incidentally contaminated *N. punctiforme* PCC 73102. In addition, the four most abundant amplicons from 16S rRNA amplicon sequencing from the *Nostoc* sp. KVJ3 and three from the *Nostoc* sp. KVJ2 microbiome were included in the tree.

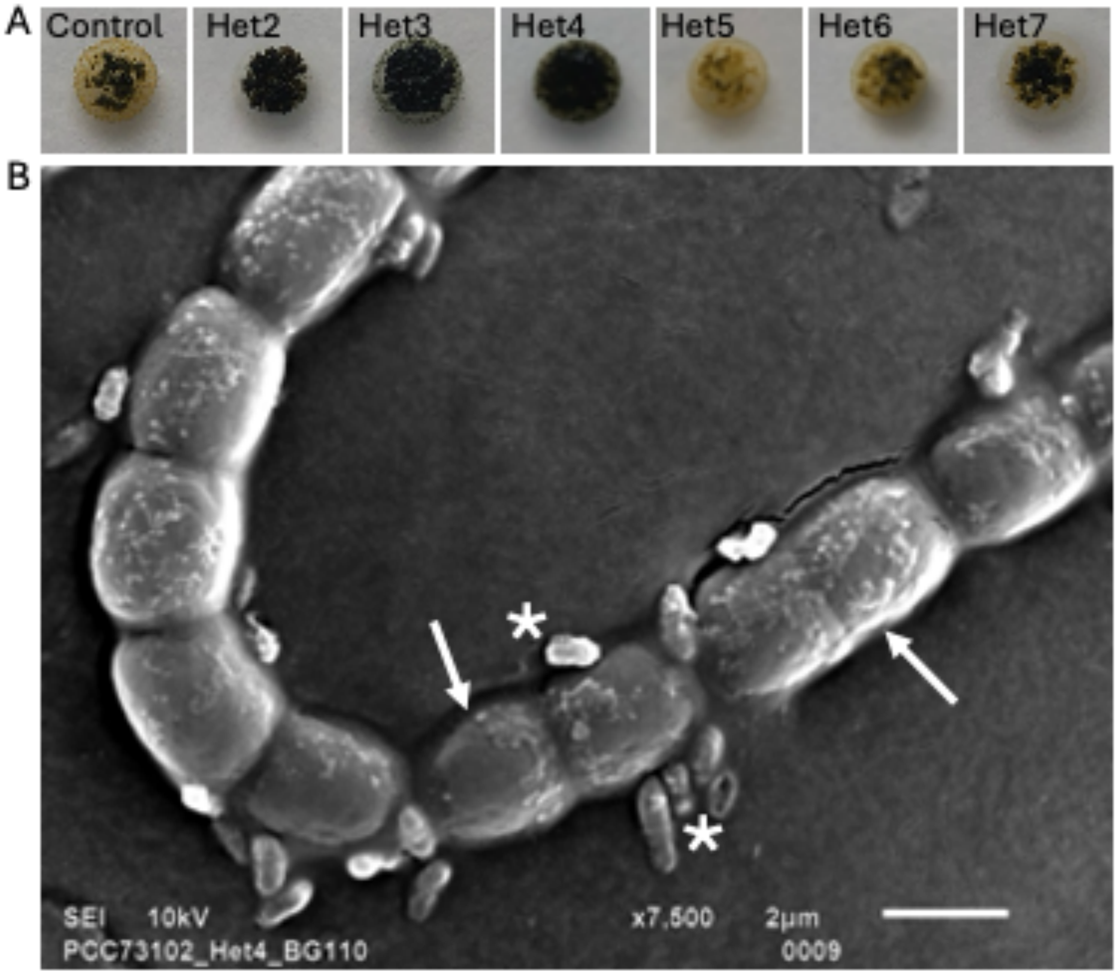


**Figure S2** Heterotrophic bacterial isolates promote growth of *N. punctiforme* PCC 73102. (A) Several isolated heterotrophic bacteria enhanced the growth of *N. punctiforme* PCC 73102 on BG11$_0$ agar plates. Particularly Het 3, 4 and 8 had a strong growth-promoting effect on the *N. punctiforme* PCC 73102. (B) Scanning electron microscopy showed an intimate interaction with *A. tumefaciens* Het4 (asterisk) and remarkable number of heterotrophic bacteria on the cell wall of cyanobacteria (arrow).

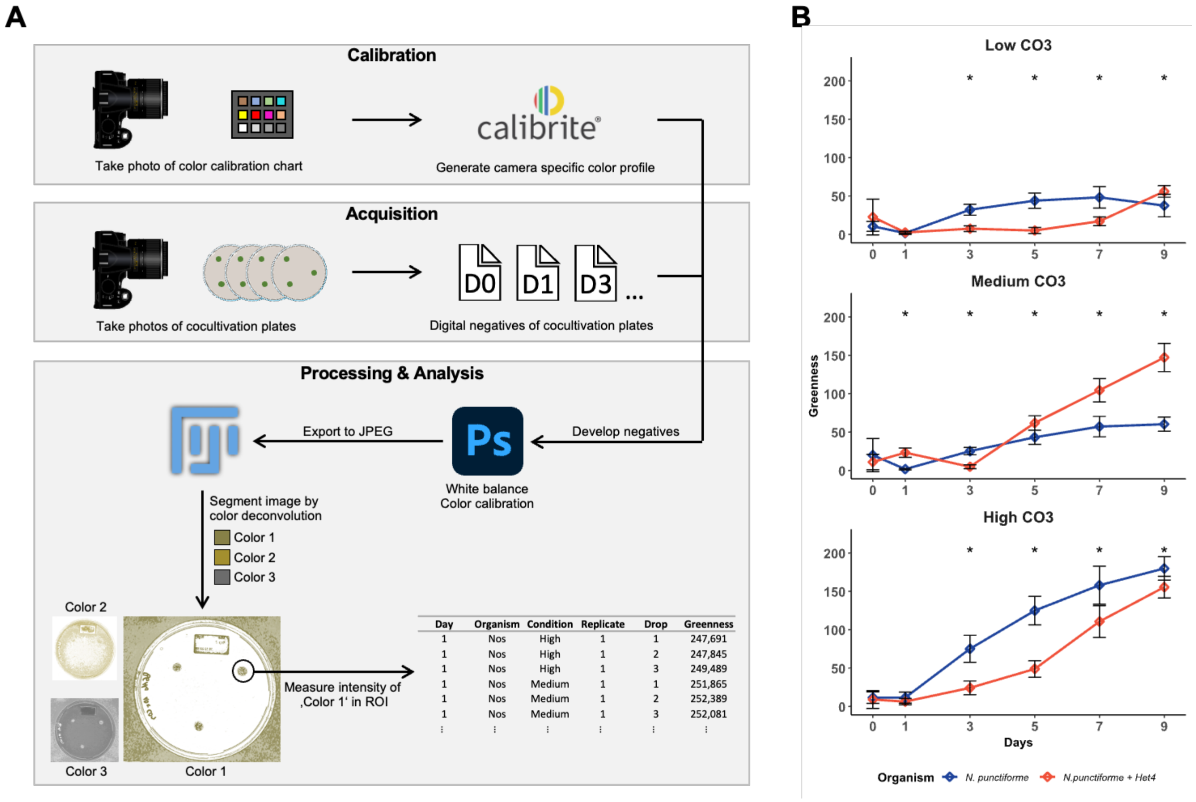


**Figure S3** Digital image analysis of growth promotion assays A) Graphical overview of the measuring method to determine Surface Greenness. In short, first a color profile for the particular camera used to take the pictures is generated using a commercially available color target and the accompanying software. Then, this color profile is used to color correct the RAW photos taken of the cultivation plates while also performing white-balancing. From a representative plate picture the values for color deconvolution and the measuring area are recorded using their respective UIs in Fiji. These values are subsequently used on every plate picture to generate the table of measurements. Data visualization is done in RStudio. Image sources: The Calibrite Logo is a registered trademark of Calibrite LLC, Wilmington, USA. petri-dish-top-gray icon by Servier https://smart.servier.com/ (CC-BY 4.0). (B) Surface Greenness measurements of axenic *N. punctiforme* PCC 73102 and *N. punctiforme* PCC 73102 + *A. tumefaciens* Het4 co-cultivation plates und various carbonate conditions. The droplet plates were prepared and incubated as previously described. Measured Surface Greenness in time points marked with an asterisk is statistically significantly different, as determined by *Welchs Two Sample t-test* using a $p < 0.05$ cutoff. T-test results are shown in Table S2.

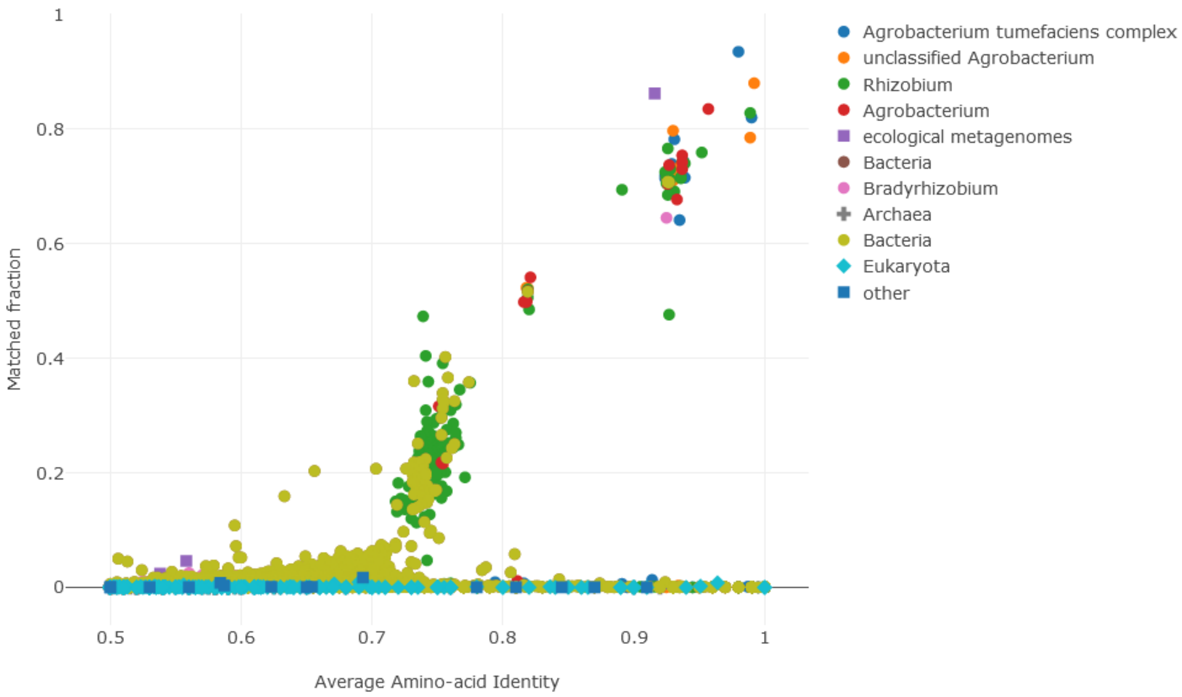


**Figure S4** Amino acid identity (AAI) of *A. tumefaciens* Het4 illustrated by AAI profiler (Medlar *et al*, 2018). AAI is plotted on the horizontal axis and the vertical axis represents the matched fraction of query proteins to the database. AAI values above 95% correspond to the same species.

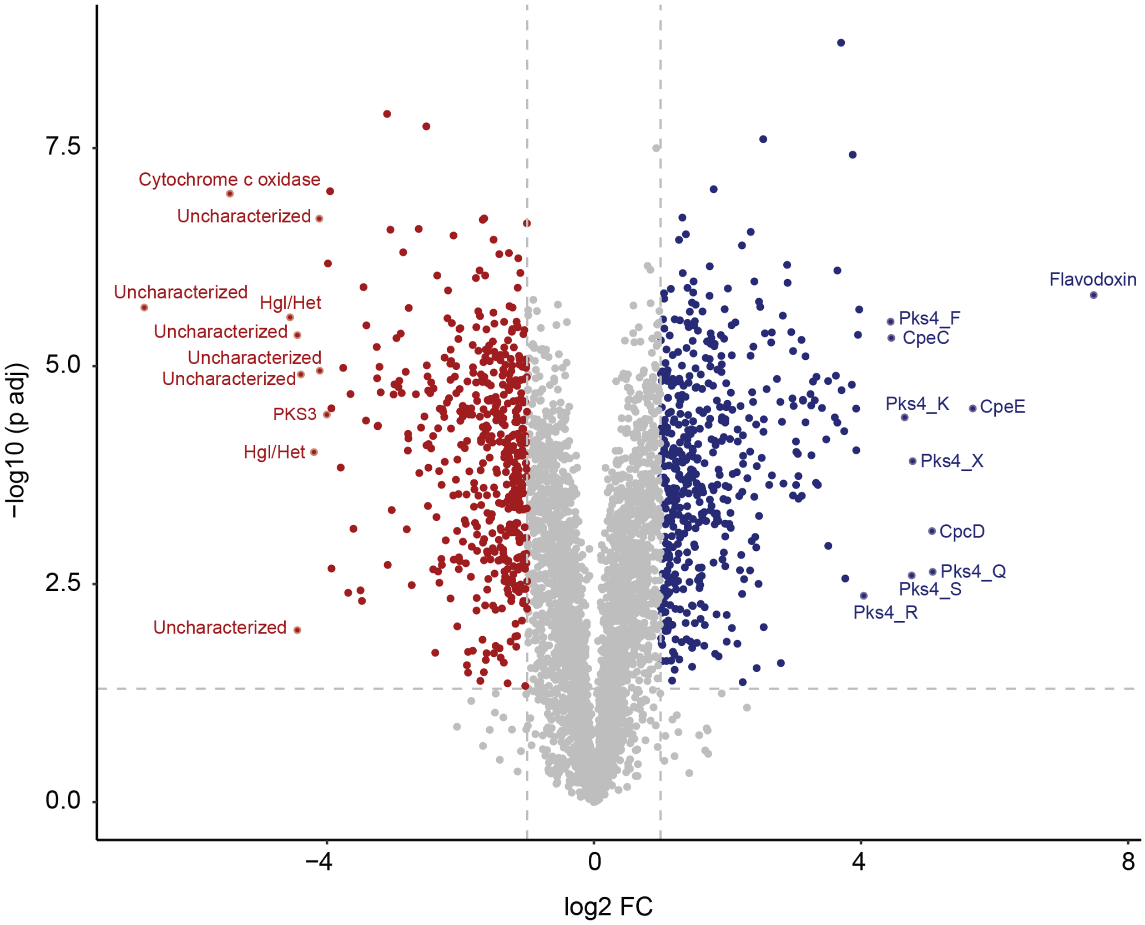


**Figure S5** Volcano plot illustrating differential expression of all proteins identified in the *N. punctiforme* PCC 73102 endoproteome in monoculture and co-culture with *A. tumefaciens* Het4. Each dot on the plot represents an individual protein and labels indicate protein name, when available. A Student's two-sample unpaired t-test with permutation-based multiple test correction was used for differential protein analysis. The fold-change and significance thresholds are indicated by vertical and horizontal dashed lines, respectively. Color coding indicates significant changes (red for downregulated, blue for upregulated, and grey for non-significant changes).

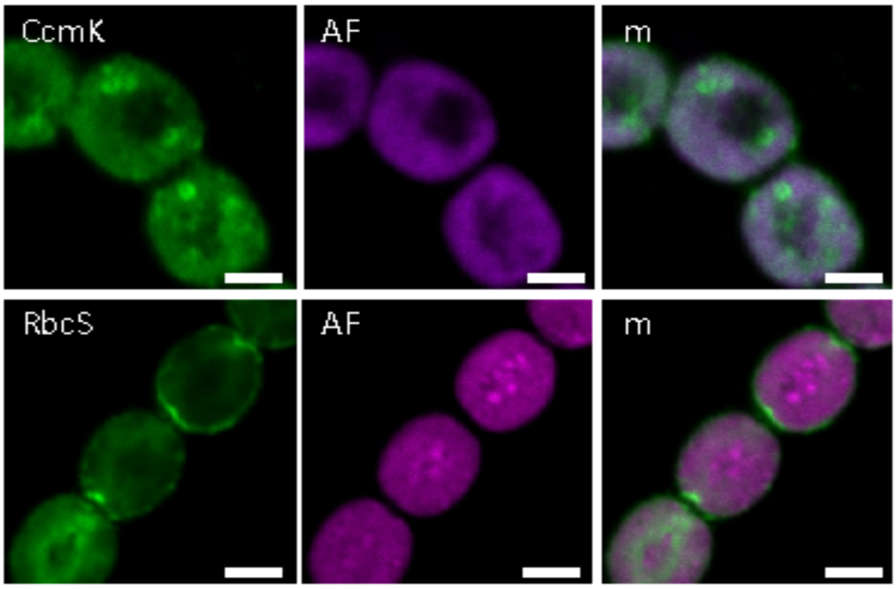


**Figure S6** Immunofluorescence labelling of carboxysomal shell protein CcmK and the small subunit of RubisCO RbcS in *N. punctiforme* PCC 73102. Immunofluorescence labeling for CcmK and RbcS in *N. punctiforme* PCC 73102. Carboxysome shell protein (CcmK2) was used to serve as a marker protein to map the placement of the carboxysomes in the cells. Typical carboxysomal ring-like structure also confirmed the integrity of the filaments in the specimen. RbcS was more evenly distributed within the cytoplasm than CcmK2 and was occasionally detected underneath the cell membrane. AF = autofluorescence, m = merged images. Scale bar 1 µm.

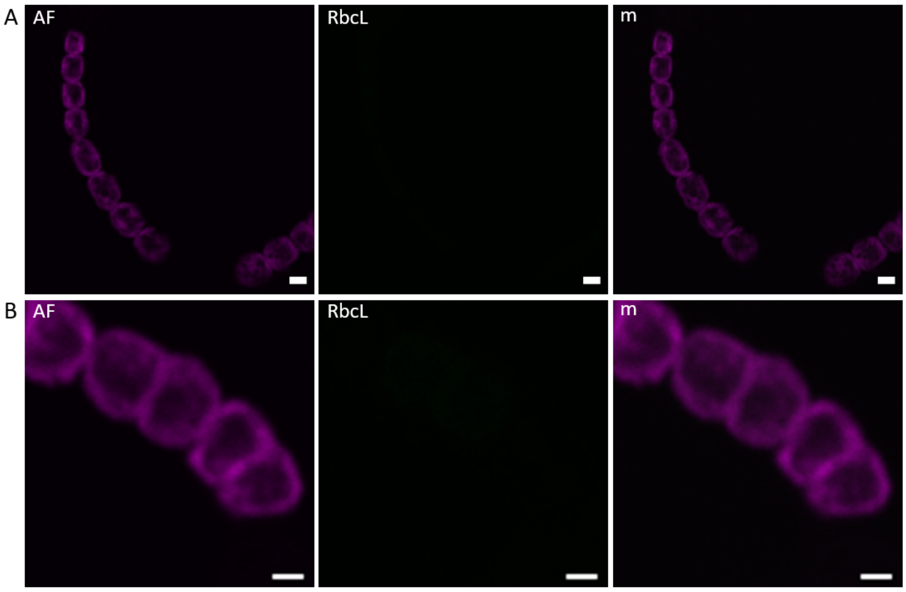


**Figure S7** Negative controls for IFM. Images were acquired by using the same parameters as in Fig. 3 and Fig. S6. Images show no binding of the secondary antibody in axenic *N. punctiforme* PCC 73102 grown on $BG11_0$. AF = phycobilisome autofluorescence, m = merged. (A) Scale bar 2 µm (B) Scale bar 1 µm.

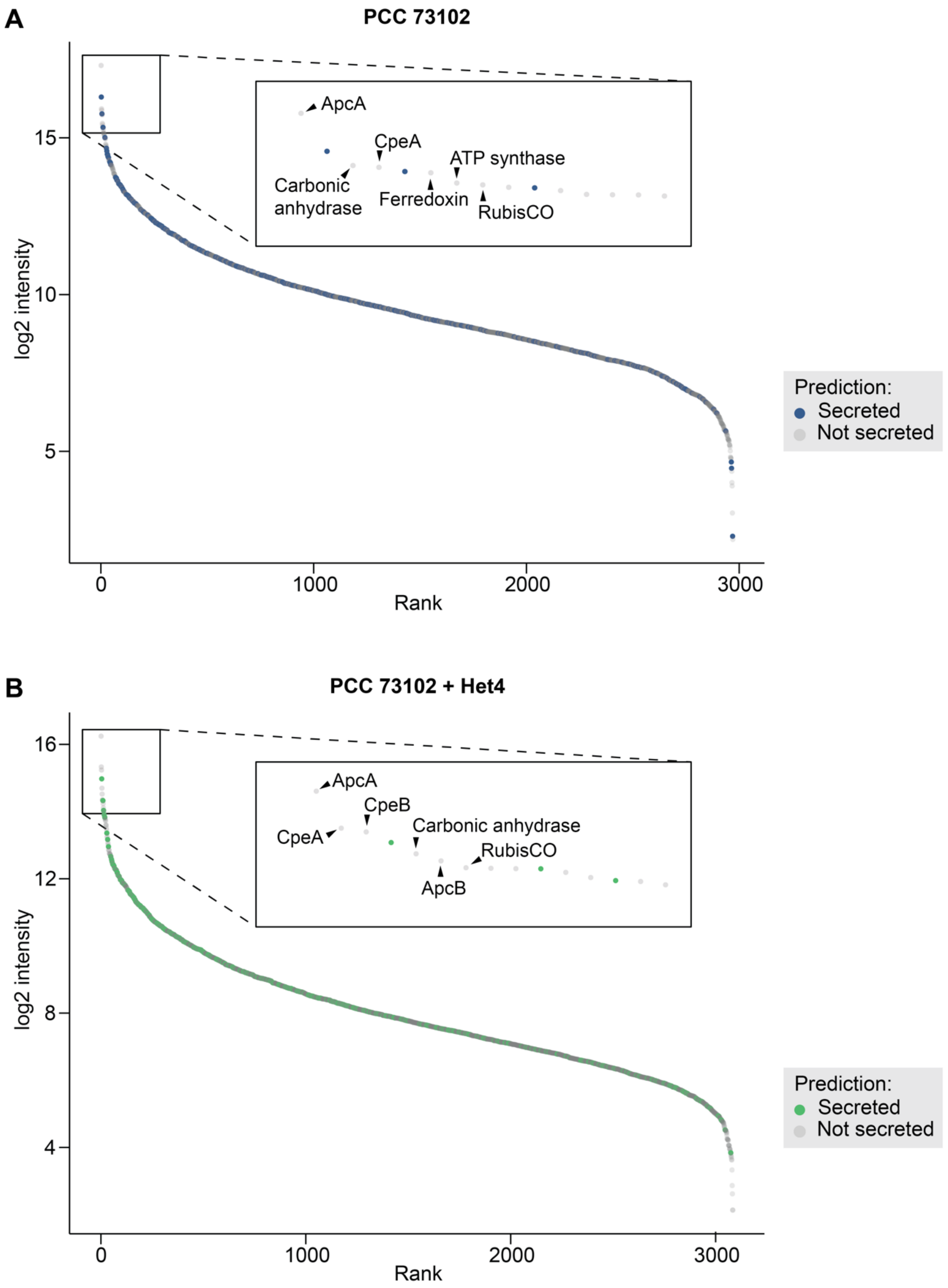


**Figure S8** Intensity of proteins identified in the *N. punctiforme* PCC 73102 exoproteome ordered by rank. The color indicates whether a protein is predicted to be actively secreted (secreted, blue (A)/ green dots (B)) or present via alternative routes (not secreted, grey dots). Panel A represents the exoproteome of *N. punctiforme* PCC 73102 in monoculture. Panel B represents the exoproteome of *N. punctiforme* PCC 73102 in co-culture with *A. tumefaciens* Het4. Secretion prediction was done according to (Russo, 2024). Dots represent means of biological replicates. Insets show the top 15 most abundant proteins identified in the exoproteome with labels indicating the 6 most abundant proteins predicted as not secreted.

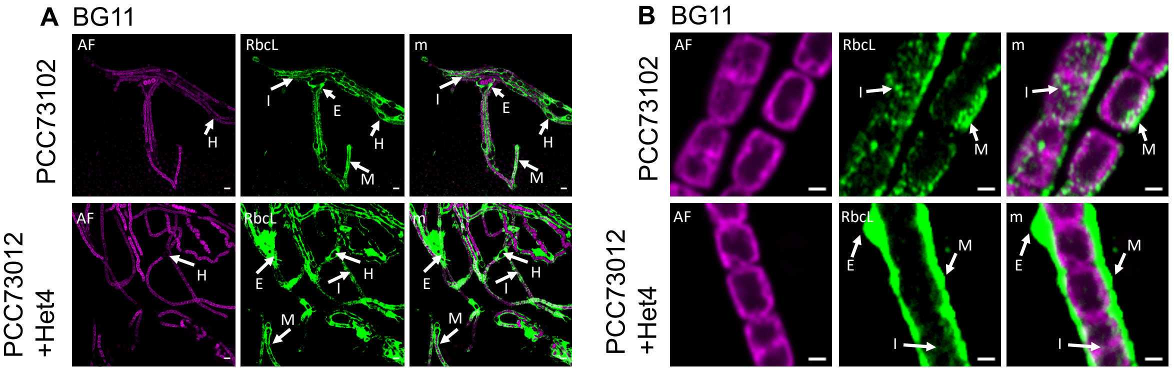


**Figure S9** Immunofluorescence detection of RbcL under nitrogen replete conditions in BG11 medium (A) Overview micrographs illustrating the phenotypic heterogeneity among filaments under nitrogen replete conditions. (B) Selected detail images showing subcellular localization of RbcL in mono-and co-cultures.

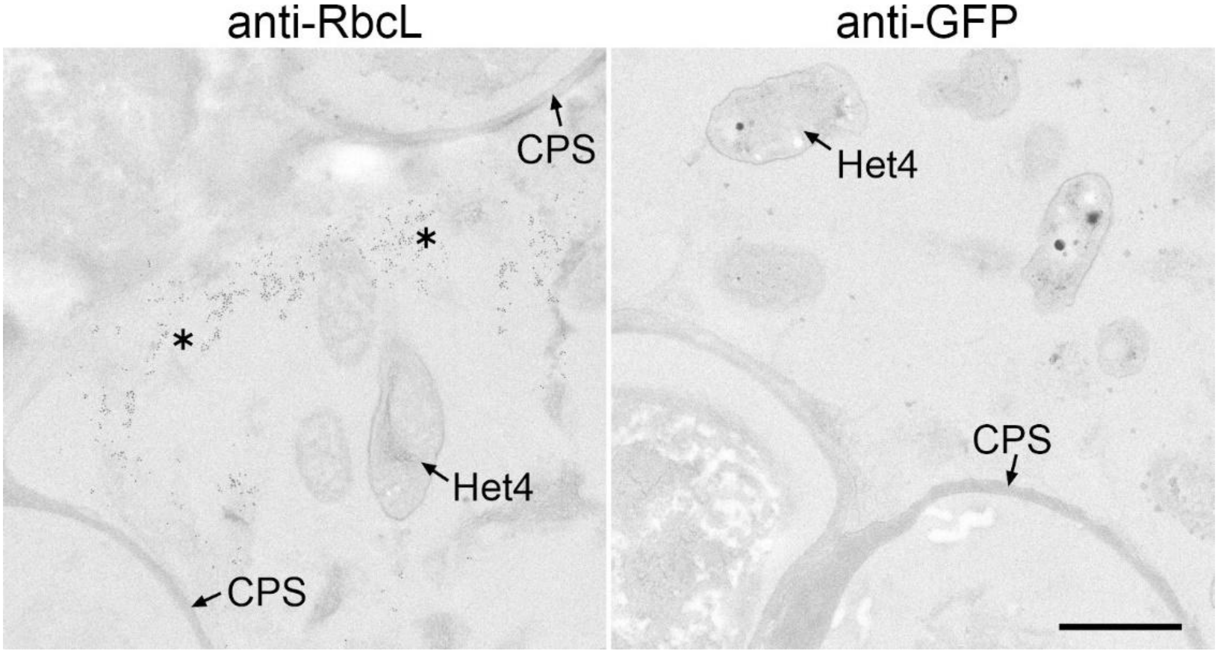


**Figure S10** Immunogold TEM. Co-culture of *N. punctiforme* PCC730102 and *A. tumefaciens* Het4 was incubated with anti-RbcL and, for control, anti-GFP. Note the numerous immunogold particles (asterisks) at the periphery of the polysaccharide capsid (CPS) in the specimen incubated with anti-RbcL. Lack of immunogold in the control specimen demonstrates specificity of anti-RbcL labeling. Bar, 1 µm
